# Api-TRACE: A System for Honey Bee Tracking in a Constrained Environment to Study Bee Learning Process and the Effect of Lithium on Learning

**DOI:** 10.1101/2024.09.24.614513

**Authors:** Babur Erdem, Ayben Ince, Sedat Sevin, Okan Can Arslan, Ayse Gul Gozen, Tugrul Giray, Hande Alemdar

## Abstract

Learning is an adaptive behavior that improves the performance of bees in foraging, dance communication, predator avoidance, and other tasks. Any deficiencies in learning could be detrimental to the long-term survival of the bee colony. Passive avoidance task is a fundamental procedure for investigating learning. We introduce Api-TRACE, a computer vision-aided system to analyze the avoidance assays. Api-TRACE tracks individual bees from the video footage of the assay and detects the moments when they were exposed to a stimulus. The algorithm provides stimulus exposure duration and learning profiles of each individual bee, enabling fast and detailed analysis of the results. Electric shock avoidance assay is one of the most common experimental methods to assess learning. We designed an apparatus for the electric shock avoidance experiments using within-reach hardware and 3D-printed components. We used Api-TRACE and experimental apparatus to investigate the effect of lithium on bee learning success in passive avoidance and reversal learning paradigms through an electric shock avoidance assay. Lithium is a potential chemical for combating *Varroa*, a bee (*Apis*) parasite, and a well-known medication for treating bipolar disorder. It has been known that lithium alters learning in humans and other animals. Before the experiment, we treated the bees with a sucrose solution with 0, 5, 25, and 125 mM LiCl *ad libitum*. Our results indicated that a decrement in learning performance emerged with increasing lithium doses in the reversal phase but not in the acquisition of the electric shock avoidance assay.

## 1. Introduction

Honey bees are crucial in agriculture and essential pollinators that significantly enhance crop yields and biodiversity. Determining learning success is essential because associative learning is key in bee foraging behavior, dance communication, and predator avoidance (Hammer & Menzel, 1995; Ings & Chittka, 2008). Learning-related behaviors are vital not only for the individual but also for the colony (Menzel & Müller, 1996). The passive avoidance task is a behavioral experiment that measures an organism’s ability to learn to avoid a negative stimulus. The avoidance learning task has been used in various studies on honey bees (Abramson, 1986; Agarwal et al., 2011; Dinges et al., 2013; Avalos et al., 2017). Electric shock avoidance assay (ESAA) is a fundamental experiment method for passive avoidance tasks. The assay has been used to investigate the effect of environmental conditions and pesticide applications on learning.

Furthermore, through the assay learning pathways have been clarified. To quantify the results of this assay, researchers view experiment videos and record the time the bees stay in the shock area individually. In other words, this quantification process takes time per observer equal to the number of bees in samples multiplied by the duration of the experiment. That becomes highly labor-intensive and introduces possible errors due to observer fatigue. In order to automatize the process, we created Api-TRACE: Honey Bee Tracking in <u>C</u>onstrained <u>E</u>nvironments to analyze ESAA videos. The Api-TRACE is a computer vision (CV) aided system to track honey bees in a constrained environment. This novel system eliminates labor and possible measurement errors.

Prior research on bee behavior and bee health employed computer-assisted techniques. CV methods were used to determine foraging activity (Ngo et al., 2019; Ngo et al., 2021), behavioral patterns (Blut et al., 2017; Kabra et al., 2013; Bozek et al., 2021), social behavior changes under an infection (Geffre et al., 2020) and nursing behavior affected by neonicotinoid pesticides (Siefert et al., 2020). Thus, the CV algorithms have been a valuable help for the researchers. The Api-TRACE system was suitable, especially for analyzing avoidance learning assays in bees and other animals (Tsaltas et al., 2007; Richter-Levin et al., 1992; Cappeliez & Moore, 1988; Xia et al., 1997; Hines & Poling, 1984; Agarwal et al., 2011; Avalos et al., 2017; Avalos et al., 2021).

In addition, the production of the experiment apparatus has limited the widespread application of ESAAs because it requires professional help. To simplify this, we designed an electric shock apparatus with rods and nuts that can easily be obtained from any hardware store or online shopping site and created 3D printing models for the plastic components. As a result, this learning test can now be easily implemented in related laboratories.

We tested our easy-to-produce experimental setup and our new software to evaluate the effect of lithium on learning in bees. The use of lithium in the fight against *Varroa*, a harmful ectoparasite affecting honey bees, began to be considered (Ziegelmann et al., 2018). However, lithium treatment has multifaceted effects on bee health. For example, while lithium reduced viral load and mitigated oxidative stress (Jovanovic et al., 2022), it caused toxicity (Kolics et al., 2021a; Kolics et al., 2021b) and significant brood damage (Rein et al., 2022). Moreover, lithium treatment affected behaviors such as movement (Hurst et al., 2014), locomotor activity, and circadian rhythms (Sevin et al., 2022; Erdem et al., 2023). Incidentally, in humans, lithium treats bipolar disorder, marked by mood shifts, increased activity, distractibility, and heightened goal-directed behavior (Alda, 2015; American Psychiatric Association, 2022). Lithium also affects learning. In humans, lithium impairs memory in free-recall tasks (Kroph & Müller-Oerlinghausen, 1979), visual-motor function, and processing speed (Judd et al., 1977). In rodents, lithium causes cognitive deficits (Wu et al., 2001), memory issues (Vasconcellos et al., 2003), and learning impairments (Richter-Levin et al., 1992). Furthermore, it was recorded that lithium impaired passive-avoidance learning in rats (Cappeliez & Moore, 1988; Hines & Poling, 1984). On the contrary, another study indicated lithium enhanced long-term retention in passive-avoidance tests (Tsaltas et al., 2007). Concerning insects, in the fruit flies (*Drosophila melanogaster*), lithium abolished memory in avoidance learning tasks (Xia et al., 1997). We found it appropriate to study the effect of lithium on bee learning because lithium may be used in hives against mites.

Our study investigates whether lithium affects the avoidance learning and reversal learning success of honey bees. The reversal learning paradigm allows the measurement of adaptive behavior and cognitive flexibility (Izquierdo et al., 2017; Claudio et al., 2018) because plasticity in learning is crucial for adapting to a constantly changing environment (Seeley, 1994; Ferguson et al., 2001). We hypothesize that lithium negatively affects the avoidance learning or its plasticity in honey bees. To test our hypothesis, we measure the learning performance in the acquisition and reversal phase in the ESAA, which has also been used in several studies previously (Avalos et al., 2021; Dinges et al., 2013).

## 2. Materials and Methods

### 2.1. Experiment setup

The experiment setup consisted of a photo studio shooting tent (diffusion softbox), laboratory stand, flat screen, webcam (mobile phone camera or any video recorder may be used), 6 V DC adapter, and electric shock apparatus. Our electric shock apparatus was designed based on the apparatus used by Dinges et al. (2013). 25 M3 threaded rods, 48 M3 nuts, two meters of copper wire, transparent acetate papers, threaded rod holders, and the shuttle boxes were used to produce the electric shock apparatus.

The production stages of the electric shock apparatus was as follows (Fig. 1):

1. Print rod holders and shuttle boxes (The download link of the 3D models was shared in the Data Availability part).
2. Cut the rods using side cutters according to the screen size.
3. Insert the rods into the rod holders.
4. Place the nuts on the end of the rods, skipping the rods one by one. Do not place any nut on the middlemost rod. Make sure only one end of each rod is nutted.
5. Cut the copper wire into four half-meter pieces. Wrap it around the ends of the nutted rods. Ensure that the copper wire does not come into contact with rods that do not have nuts on the end.
6. Place another set of nuts on the ends of the nutted rods and turn the nuts so that the copper wire is tightly sandwiched between the two nuts.
7. Cut the acetate paper to 17 x 170 mm and insert it into the slide of the shuttle box.
8. Apply a thin layer of Vaseline to both the inside surface of the shuttle box and the detachable acetate paper roof. (Vaseline prevents bees from climbing to the walls and losing contact with the electric grid).

**Fig. 1.**
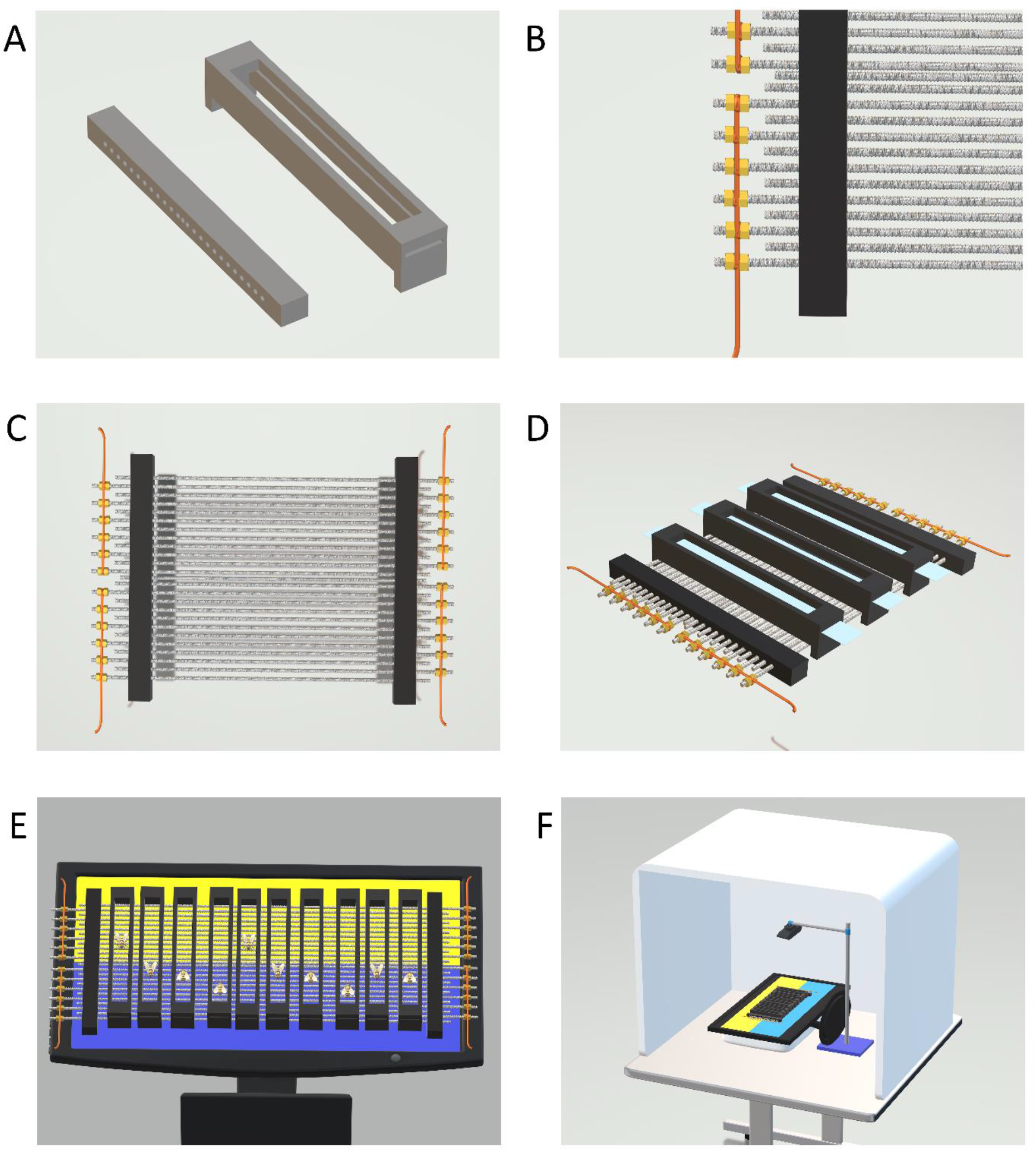
3D-printed components for the electric shock apparatus, including rod holders and shuttle boxes (A). Close-up view of the rod and copper wire assembly to build the electric grid (B). Top view of the assembled electric shock grid (C). Perspective view of the completed electric shock apparatus with shuttle boxes aligned on the grid (D). The electric shock apparatus was placed on the flat screen display showing the delineation of shock and safe areas (E). The fully assembled experiment setup was placed inside a photo studio shooting tent to ensure homogeneous lighting conditions (F).

Configuration of experiment setup (Download link of the 3D illustrations of experiment setup was shared in the Data Availability part):

1. Mount the webcam to the laboratory stand.
2. Place the flat screen parallel to the floor.
3. Move the lab stand close to the screen. The camera should see the entire screen surface. It should be centered and perpendicular to the screen.
4. Put the electric shock apparatus on the screen. The middlemost rod of the shock grid should align with the separation of the colors representing the shock and safe area reflected on the screen.
5. Line up the shuttle boxes on the electrical grid.
6. The experiment setup should be placed inside the photo studio shooting tent (diffusion softbox) to ensure homogeneous light conditions and prevent reflections and shadows.
7. Connect the positive and negative outputs of the adapter to the copper wires on the side of the electrical grid. A DC jack to alligator clip power adapter cable may also be used.
8. Transfer the bees into the shuttle boxes. Each shuttle box should contain a single bee that could freely traverse between the shock and safe areas. Then, plug the adapter into the socket.

### 2.2. Api-TRACE Video Processing Module

The video processing module (VPM) was developed with Python (>=3.6) to track the movement of bees in video footage and analyze whether they were exposed to the stimulus, which was the electric shock in our experiments, during a specific time interval. We used CV techniques to track the bees’ positions and did geometric calculations to determine whether the bees were within a defined exposure area (Fig. 2). We used “*OpenCV”,* “*NumPy*”, “*Shapely*”, and “*MoviePy*” libraries. In order to achieve accurate tracking and analysis, we used the following stages:

**Fig. 2.**
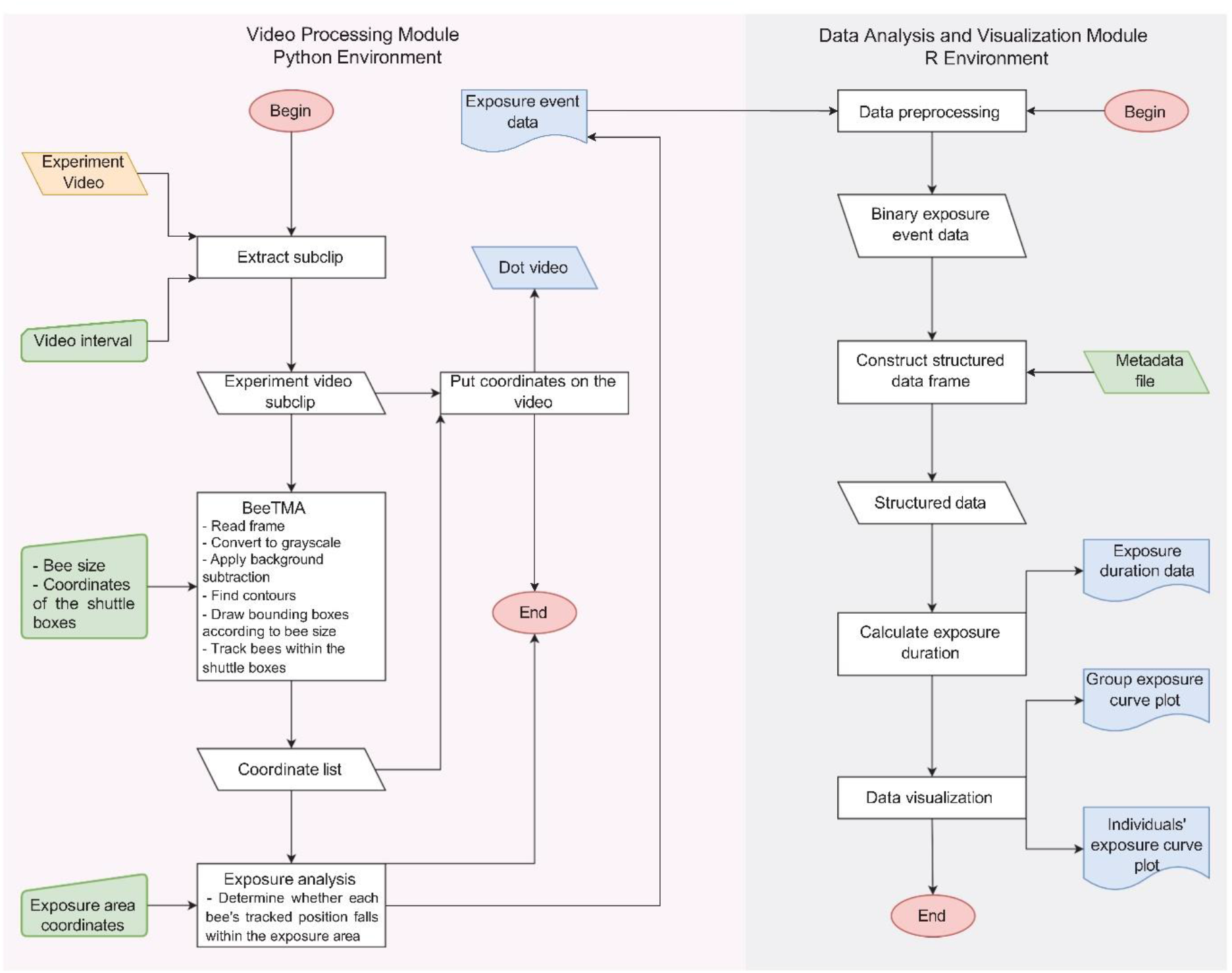
Flowchart of video processing module and data analysis and visualization module.

#### Video Processing

The recorded video was processed to extract a subclip containing the desired trail of the experiment. The *ffmpeg_extract_subclip* function from the “*moviepy*” library was employed. The subclip spanned from the stimulus initiation time to a defined duration of the trial of the experiment.

#### Bee Size Measurement

The user was prompted to measure the length of a bee in the video. The program allowed the user to draw a line on the first frame of the cut video frame via a graphical user interface (GUI) using the *OpenCV* library to measure the bee’s size (Fig. 3A). The minimum and maximum sizes of a bee were calculated based on the user’s measurement. These measurements were then used in the “Bee Tracking and Motion Analysis” stage for thresholding and dilation operations.

**Fig 3.**
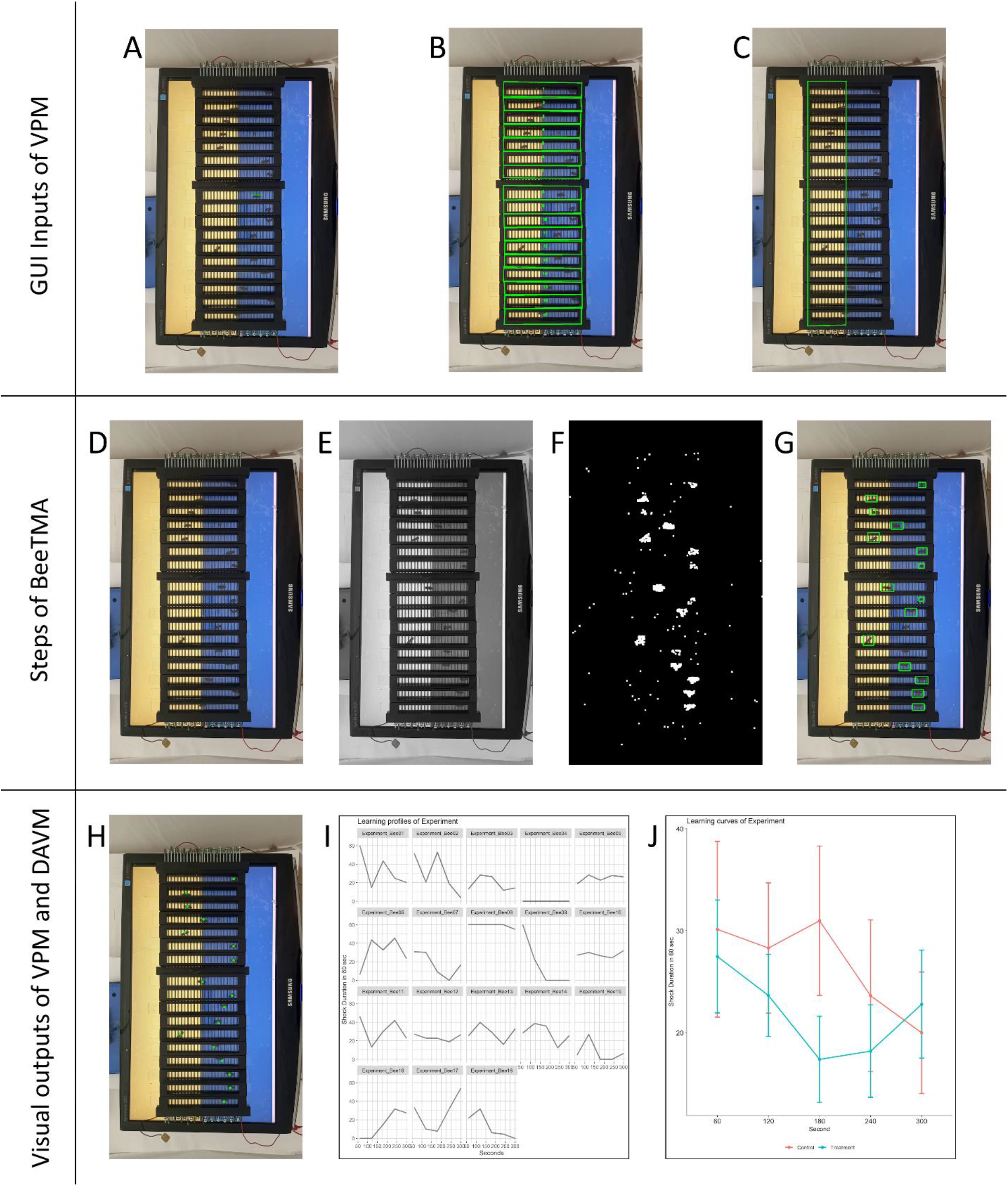
Images of the video processing module and data analysis and visualization module.

#### Defining Regions of Interest (ROIs)

Two types of ROIs were defined on the video frames for subsequent analysis:

1. Shuttle Boxes: Areas of interest where bee movement was tracked. These were manually defined by drawing polygons around specific regions of the video frame via GUI. The polygons served as borders of the shuttle boxes to track bee movement within the bordered area (Fig. 3B).
2. Exposure Area: The specific region where exposure event occurred. This area was manually drawn as defined for the shuttle boxes by drawing a polygon on the video frame via GUI (Fig. 3C).

Bee Tracking and Motion Analysis Algorithm (BeeTMA):

Bee movement was tracked and analyzed within the defined shuttle boxes. The process involved the following steps:

1. A frame was read (Fig. 3D) and converted to grayscale (Fig. 3E).
2. Background subtraction was performed using the Gaussian Mixture Model algorithm (*cv2.createBackgroundSubtractorMOG2*) to identify the region of motion in the video frames.
3. The motion mask obtained from the background subtraction was processed using thresholding and dilation operations to extract motion areas.
4. Contours were identified within the motion areas using the *cv2.findContours* function.

Bees were identified based on their size. In the beginning, proper minimum and maximum sizes were assigned as variables in pixels (Fig. 3F).

1. Bounding boxes were drawn around the detected bees, and their center coordinates were recorded for further analysis (Fig. 3G).

Detection of the Bees Exposed to Stimulus:

To determine whether individual bees were exposed to a stimulus, the “*Shapely*” library is used for geometric calculations. The positional information of bees within the exposure area was compared against the defined exposure polygon. Bees whose center coordinates fell within the exposure area were considered as receiving the stimulus.

#### Data Output and Visualization

The following outputs were generated for analysis and visualization purposes:

1. A text file was created to record exposure event data. Each row of the file represented a video frame, and columns indicated whether each bee was exposed to the stimulus (True) or not exposed (False) during that time frame.
2. A video was generated, displaying the tracked positions of the bees within the shuttle boxes throughout the experiment. Each bee’s position was represented by a single dot (Fig. 3H).

A video tutorial on using the algorithm is provided as supplementary material (Video S1). The download link of the VPM script was shared in the Data Availability part.

### 2.3. Api-TRACE Data Analysis and Visualization Module

In the Data Analysis and Visualization Module (DAVM), we provided R scripts to facilitate the statistical analysis of the exposure event data obtained from the VPM. DAVM served to calculate exposure durations, export the refined data to a tab-delimited text file for further analysis, and generate accurate visualizations (Fig. 2). This module commenced by loading essential R packages, namely “*dplyr*”, “*ggplot2*”, and “*ggpubr*”, which enabled efficient data manipulation, visualization, and exporting capabilities. It had the following steps:

#### Data Preprocessing

The script read the exposure event data. The categorical exposure responses (“True” and “False”) were transformed into a numeric format (1 for “True” and 0 for “False”). This conversion laid the foundation for subsequent calculations.

#### Construction of Structured Data Frame

A new exposure response data frame was created to structure the processed data. This data frame was designed to store essential information. The data frame included Bee ID, Time, and Exposure Duration. Also, it contained independent variables and experiment-related information, such as treatment, dose, subspecies, phase, and replicate, which came from the metadata file.

#### Each column was populated with relevant details to enable organized analysis. Calculation of Exposure Duration

Exposure durations were calculated by summing exposure responses over defined time intervals. The cumulative sum of exposure responses was computed by iterating through the exposure response data, providing a quantitative representation of exposure duration for each interval.

#### Data Export

The processed exposure duration data was exported to a tab-delimited text file. These refined data were then used for further statistical analysis.

#### Data Visualization

Two types of plots were generated to represent the exposure duration patterns visually. The first type involved generating line plots for individual bees, illustrating each bee’s exposure duration over time (Fig. 3I). The second type employed grouped line plots to compare exposure duration trends across independent variables (Fig. 3J). These plots showcased the mean and standard error of exposure duration over time, aiding in identifying potential treatment effects.

The download link of the DAVM script was shared in the Data Availability part.

### 2.4. Lithium experiments

The honey bee colonies used in this study were obtained from the apiary of Ankara University’s Veterinary Faculty. The hive entrances were obstructed by using plastic wire mesh to capture returning forager bees. The bees that accumulated on the mesh were collected into small cages. The samples were collected from at least three distinct hives and pooled for each subsequent analysis to mitigate potential colony variations.

Collected forager bees were individually placed into separate hoarding cages. Each cage received sustenance from an overhead syringe filled with a 50% (w/v) sucrose solution containing a predetermined quantity of LiCl. Four experimental groups were formed: the low-dose group (5 mM LiCl), the medium-dose group (25 mM LiCl), the high-dose group (125 mM LiCl), and the control group (sucrose-only). We determined the doses by taking the square and square root of 25 mM, used in previous studies (Ziegelmann et al., 2018; Kolics et al., 2021b; Rein et al., 2022). The feeding regimen was upheld overnight for a total of 16 hours. The hoarding cages were kept in an incubator at a constant temperature of 33 °C and a relative humidity of 65%, all within a controlled dark environment. The experiment groups were then subjected to the ESAA.

An LG Flatron W2243S computer monitor displayed one side as blue (hex code: #376092) and the other as yellow (hex code: #FFFF00). A Logitech C920 webcam captured experiment video records, and a Godox DF-01 portable diffusion box (70×70×70 cm) was used to place the experimental setup (Fig. S1).

Experiments were conducted under ambient light and room temperature. The experimental procedure was as follows:

1. Habituation: Bees were introduced to the experimental setup, and the computer monitor was turned off for an initial habituation period of 10 minutes.
2. Acquisition phase: One of the colors was designated as the shock side (CS+), and the electric current was channeled to the CS+ side for 5 minutes.
3. Interval between phases: The electric current was deactivated, and the monitor was closed to allow the bees to rest for 10 minutes.
4. Reversal phase: The shock sides were switched, with the opposite color as the new CS+. This reversal phase also persisted for 5 minutes.

To ensure counterbalancing, the color assignments of CS+ and CS-were reversed for subsequent batches of bees. Multiple batches of bees were analyzed, with each group comprising at least two counterbalanced batches. The number of bees at the beginning of the experiment was 33, 36, 36, and 32 in the control, low-dose, medium-dose, and high-dose groups, respectively. If a bee was completely motionless throughout the experiment or the tracking code incorrectly pointed to the bee, those bees were not included in the statistical analysis. As a result of these eliminations, the sample sizes were 27, 33, 31, and 28 in the control, low-dose, medium-dose, and high-dose groups, respectively.

All statistical analyses were conducted in RStudio with the R 4.3 version (R Core Team, 2020). The normality of the sample distributions was checked with the Shapiro-Wilk test. We evaluated the electric shock experiment results with a linear mixed-effects model (LMEM), which allows within-group errors, using the “*lme*” function from the “*nlme*” package. In addition, the Spearman correlation test was used to determine the association between the shock duration and increasing doses of lithium for the acquisition and reversal phases.

## 3. Results

Our BeeTMA algorithm processed 16 videos of the experiment to track bee motion in 274 ROIs that define shuttle boxes. The output videos displaying the bees’ tracked positions were visually examined. The visual examinations indicated that the algorithm could not track the bees in only 8 ROIs. Thus, the algorithm’s performance was 97.08% in terms of successfully tracked ROIs.

Moreover, we determined the run time of our BeeTMA algorithm on computers with different configurations. We used an experiment video (720 p and 30 fps) that included 18 bees and lasted 5 minutes. The average processing time we obtained from the randomly selected computers was 8.20 minutes, the slowest processing time was 15.60 minutes, and the fastest was 3.61 minutes (Table 1). If an observer examined this video, the observer would have to spend 5 minutes on each bee, and the time spent on this video would be 90 minutes.

**Table 1.** Video processing time of our computer vision algorithm on computers with different configurations.

| CPU | Core number | Freq (GHz) | RAM (GB) | Avg. Duration (sec) |
| --- | --- | --- | --- | --- |
| AMD Ryzen 3 3100 | 4 | 3.60 | 16 | 429.31 |
| AMD Ryzen 7 5800HS | 8 | 2.80 | 16 | 449.26 |
| Apple M1 | 8 | 3.20 | 16 | 452.12 |
| Intel i3 1005G1 | 2 | 1.20 | 8 | 460.95 |
| Intel i5 7200U | 2 | 2.50 | 4 | 935.76 |
| Intel i7 11800H | 8 | 2.30 | 16 | 216.55 |
| Intel i7 1255U | 2 | 1.70 | 32 | 499.32 |

Then, we determined lithium’s effect on bees’ learning using our algorithms. In the acquisition phase, significant results were found in the time effect (*β* = -0.015, *F* (1, 474) = 10.721, *p* = 0.001). However, the group effect (*β* = -0.011, *F* (1, 117) = 0.324, *p* = 0.570) and interaction effect (*β* < 0.001, *F* (1, 474) = 2.128, *p* = 0.145) were not significant according to LMEM (Fig. 4A). Then, we also applied a permutation test to our model because the distribution was not normal according to the Shapiro-Wilk test (*p* < 0.05). The permutation test confirmed the significant LMEM results for the time effect (*B* = 1000, *p* = 0.001). The results suggested that in the acquisition phase, all groups learned to avoid the color associated with electric shock over the 5-minute. Thus, lithium did not affect learning in the acquisition phase. In the reversal phase, we did not find significant results in the time effect (*β* = -0.003, *F* (1, 474) = 0.353, *p* =0.553), group effect (*β* = -0.020, *F* (1, 117) = 0.956, *p* = 0.330), and interaction effect (*β* < 0.001, *F* (1, 474) = 0.038, *p* = 0.845) according to LMEM (Fig. 4B).

**Fig. 4.**
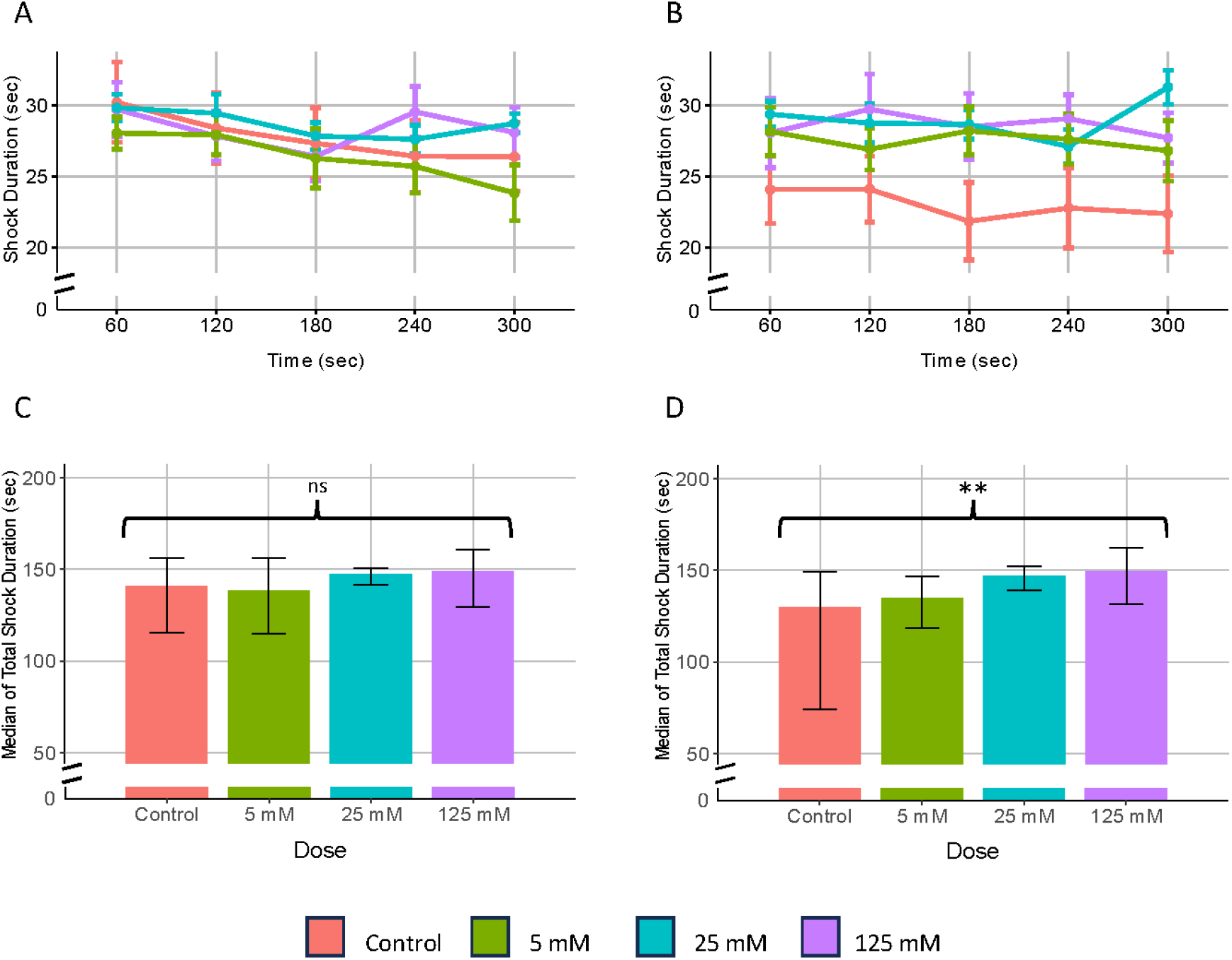
Comparison of spatial-avoidance learning rate across the groups during the acquisition phase (A) and reversal phase (B) of ESAA. Each data point shows the mean (± standard error) of the time honey bees spent on the shock side in one minute. The bars represent the median of the total duration of exposure to electric shock groups during the acquisition phase (C) and reversal phase (D). Error bars showing interquartile range. Curly brackets were used to represent the result of the Spearman correlation to determine the relationship between the duration of exposure to electric shock and lithium doses. Asterisks indicate the level of statistical significance: *p ≤ 0.05, **p ≤ 0.01, ***p ≤ 0.001, n.s. not significant.

We calculated the median and interquartile range (Q1 – Q3) of shock duration of doses for both phases. There were 140.700 (115.333 – 156.350) for control, 138.567 (114.833 – 156.367) for low-dose, 147.200 (141.767 – 150.650) for medium-dose, and 148.950 (129.550 – 160.567) for high-dose groups in the acquisition phase. In the reversal phase, there were 129.967 (74.350 – 149.267) for control, 134.733 (118.633 – 146.667) for low-dose, 147.067 (139.267 – 152.317) for medium-dose, and 149.300 (131.625 – 162.133) for high-dose groups (Table S1). Then, we determined the relationship between the duration of exposure to electric shock and lithium doses in each phase. We used a non-parametric Spearman correlation test because the normal distribution was not met according to the Shapiro-Wilk test (*p* < .05). There was no correlation found in the acquisition phase (*r* (117) = 139, *p* = 0.131) (Fig. 4C). However, a positive correlation was found in the reversal phase (*r* (117) = 0.282, *p* = 0.002) (Fig. 4D). According to this result, it has been observed that increased lithium doses negatively affect learning only in the reversal phase.

## 4. Discussion

In this study, we developed and successfully validated Api-TRACE system to analyze the ESAA in honey bees. By automating the tracking of individual bees and the detection of stimulus exposure, we significantly reduced the labor and potential observer errors typically associated with manual analysis. Our results demonstrated the BeeTMA algorithm’s performance in terms of correctly tracked ROIs as 97.08%. In the broader sense, the ability to accurately measure and analyze learning success through automated means allows for more detailed and large-scale investigations into various behaviors.

In our study, we investigated the potential impact of lithium on honey bee learning through passive avoidance and reversal learning paradigms through an electric avoidance assay using our experimental setup and algorithms. The results of LMEM revealed a significant time effect in the acquisition phase but not in the reversal phase of the experiment. Thus, the duration of stay in the electric area decreased over time at the acquisition phase, indicating that learning could occur. However, this was not the case for the reversal phase. A positive correlation was found between lithium doses and shock duration in the reversal phase, suggesting that increased lithium doses negatively affected learning during the reversal phase.

In explaining the results, it is essential to consider previous research on lithium’s effects on avoidance learning. Lithium has exhibited both favorable (Tsaltas et al., 2007) and adverse effects (Richter-Levin et al., 1992; Cappeliez & Moore, 1988; Xia et al., 1997; Hines & Poling, 1984) on avoidance learning. These studies also showed the dual effect of lithium.

One of the molecular pathways regarding the effect of lithium on learning can be established through the relationship between lithium and GSK-3 protein. The inhibitory effect of lithium on GSK-3 (ortholog of the Shaggy found in insects) activity has been demonstrated in studies on many organisms (Klein & Melton, 1996; Stambolic et al., 1996; Hedgepeth et al., 1997; Hong et al., 1997; Williams & Harwood, 2000; Shaldubina et al., 2001; Jope, 2003; Zhan et al., 2003; Castillo-Quan et al., 2016). It was later determined that GSK-3 activation had a detrimental effect on learning, memory, signal transduction, and caused habituation (Grigor’yan, 2014; Storozheva et al., 2015; Jope et al., 2017; Franciscovich et al., 2008; Beurel et al., 2015; Wolf et al., 2007). The impairment of long-term potentiation caused by GSK-3 is recovered by lithium (Hooper et al., 2007). Thus, the positive impact of lithium on learning may be elucidated by its capacity to inhibit GSK3, given that the detrimental effects of GSK-3 activity on learning have been established.

Nevertheless, an alternative pathway should be considered to elucidate the negative consequences of lithium. In this context, the impact on dopamine pathways becomes pertinent. Lithium inhibits dopamine-mediated behaviors, such as motor activity, by interfering with GSK-3-mediated pathways in mice (Beaulieu et al., 2004). Dopamine is essential for learning olfactory-electric shock conditioning in fruit flies (Zhou et al., 2019). It is shown that in fruit flies, the unexpected uncoupling of an olfactory cue from electric shock punishment activated reward-encoding dopaminergic neurons through reduced punishment-encoding dopaminergic neuron activity in a reversal learning experiment (Davoudian & Nitabach, 2021). Also, dopamine release was documented in the brains of honey bees receiving electric shocks (Jarriault et al., 2018). It is shown that blocking dopaminergic receptors had a detrimental effect on learning in ESAA in honey bees (Vergoz et al., 2007). Lastly, dopamine inhibitors decreased learning success in a dose-dependent manner in honey bees during a spatial avoidance conditioning assay, the same passive avoidance task we used in the current study (Agarwal et al., 2011). These findings suggest that the dual nature of lithium’s effects on avoidance learning involves intricate interactions with both GSK-3 and dopamine pathways. In our case, it is possible to explain the negative effect of lithium on learning, which we observed in the reversal phase of the ESAA, by the dominance of the dopamine mechanism in reversal learning. Such dominance was exemplified in a related study (Costa et al., 2015).

An unexpected situation we encountered in our experiments was that the shock duration in the control group was lower in the reversal phase than in the acquisition phase. Initially, we predicted that learning to avoid the other color associated with the electric shock during the reversal phase would take more time. This situation can be explained via a “rule learning” concept. The term “rule learning” in the context refers to the honey bees’ ability to understand general principles or rules from their environment. Bees can go beyond basic learning and comprehend broader concepts such as categorization, contextual learning, and abstract rules (Giurfa, 2007). In this case, instead of associating a color with an electric shock and being conditioned to avoid that color permanently, the bees may have learned the rule of positioning themselves opposite the color that caused the electric shock. This interpretation suggests a higher cognitive flexibility and adaptability level in honey bee behavior.

## 5. Conclusions

Assessing learning success is crucial since associative learning underpins bee foraging, dance communication, and predator avoidance, all vital for the colony’s survival. The passive avoidance task, which measures an organism’s ability to avoid negative stimuli, is a key experiment for studying honey bee behavior. The ESAA is a fundamental method in those tasks, useful for exploring the effects of many independent variables on learning. In this study, we developed an easily constructible experimental apparatus and an algorithm to analyze the experiment videos. We then used these to determine the effects of lithium, which is recommended as an acaricide, on learning.

Our Api-TRACE system provides precise and efficient tracking of bee movement during the avoidance assay. This tool enhances the accuracy of behavioral data collection and analysis, offering a significant improvement over traditional manual observation methods.

The design of the experimental apparatus allowed us to use standard hardware and 3D-printed components. By lowering the cost and technical barriers associated with setting up the ESAA, laboratories conducting related research can adopt and implement this method, leading to a broader application of avoidance learning studies in honey bees. Moreover, the application of our algorithm and apparatus can also be extended to other organisms and experimental achived in a constrained environment. Exploring the use of our system in studying learning in other insect species or small animals could broaden the scope of behavioral research.

Notably, the results of our study suggest a negative impact of lithium on honey bee learning. If lithium affects the learning and cognitive abilities of bees, it could have detrimental consequences for the overall health and productivity of the colony. Therefore, our results emphasize the importance of careful consideration and monitoring when using lithium as an acaricide in honey bee colonies.

In conclusion, our study contributes to further understanding of how chemical exposures affect bee behavior. Also, it provides a technique and a method for future studies using various organisms. The dimensions of the setup that we used can be changed depending on the organism to be investigated. Since VPM does not include a step to identify the honey bee species specifically, it can be easily extended to various organisms.

## Acknowledgments

This article was produced from the research conducted by the corresponding author as part of his doctoral dissertation. This work was supported by The Scientific and Technological Research Council of Türkiye [TÜBİTAK grant number: 122E014]; the Middle East Technical University Research Fund [BAP grant number: ADEP-302-2024-11468]; The U.S. National Science Foundation [Award numbers: 2318597; 2231637; 1736019]; The USDA-NIFA [Award number: 2021-67014-34999]. Thanks to Tuna Ercan for his help in producing 3D parts, and Assoc. Prof. Dr. Ali Emre Turgut and Assoc. Prof. Dr. Erol Sahin for providing workspace, tools, and materials.

## Data Availability

The data of the lithium experiment, the scripts of the Api-TRACE, 3D models of the parts used in the electric shock apparatus, and 3D illustrations of the experiment setups are available in the Zenodo repository: https://doi.org/10.5281/zenodo.13375378

Also, the scripts that will include possible future updates can be accessed from this repository: https://github.com/baburerdem/Api-TRACE

## Author Statement

**Babur Erdem:** Conceptualization, Methodology, Software, Formal analysis, Investigation, Writing - Original Draft, Visualization. **Ayben Ince:** Formal analysis, Investigation. **Sedat Sevin:** Investigation, Resources. **Okan Can Arslan**: Investigation. **Ayse Gul Gozen:** Writing - Original Draft. **Tugrul Giray:** Conceptualization, Methodology, Writing - Original Draft, Funding acquisition. **Hande Alemdar:** Conceptualization, Methodology, Resources, Writing - Original Draft, Funding acquisition

## REFERENCES

1. Abramson, C.I. (1986). Aversive conditioning in honeybees (Apis mellifera). Journal of Comparative Psychology, 100(2), 108–116. 10.1037/0735-7036.100.2.108.

2. Agarwal, M., Giannoni Guzmán, M., Morales-Matos, C., Del Valle Díaz, R. A., Abramson, C. I., & Giray, T. (2011). Dopamine and octopamine influence avoidance learning of honey bees in a place preference assay. PLoS ONE, 6(9), 1–9. 10.1371/journal.pone.0025371.

3. Alda, M. (2015). Lithium in the treatment of bipolar disorder: pharmacology and pharmacogenetics. Molecular Psychiatry, 20(6), 661–670. 10.1038/mp.2015.4.

4. American Psychiatric Association. (2022). Bipolar and Related Disorders in Diagnostic and Statistical Manual of Mental Disorders: DSM-5-TR (ed. Ostacher, M. J.) 150–169. American Psychiatric Association Publishing.

5. Avalos, A., Pérez, E., Vallejo, L., Pérez, M. E., Abramson, C. I., & Giray, T. (2017). Social signals and aversive learning in honey bee drones and workers. Biology Open, 6(1), 41–49. 10.1242/bio.021543.

6. Avalos, A., Traniello, I. M., Claudio, E. P., & Giray, T. (2021). Parallel mechanisms of visual memory formation across distinct regions of the honey bee brain. Journal of Experimental Biology, 224(19). 10.1242/jeb.242292.

7. Beaulieu, J. M., Sotnikova, T. D., Yao, W. D., Kockeritz, L., Woodgett, J. R., Gainetdinov, R. R., & Caron, M. G. (2004). Lithium antagonizes dopamine-dependent behaviors mediated by an AKT/glycogen synthase kinase 3 signaling cascade. Proceedings of the National Academy of Sciences of the United States of America, 101(14), 5099–5104. 10.1073/pnas.0307921101.

8. Beurel, E., Grieco, S. F., & Jope, R. S. (2015). Glycogen synthase kinase-3 (GSK3): Regulation, actions, and diseases. Pharmacology and Therapeutics, 148, 114–131. 10.1016/j.pharmthera.2014.11.016.

9. Blut, C., Crespi, A., Mersch, D., Keller, L., Zhao, L., Kollmann, M., Schellscheidt, B., Fülber, C., & Beye, M. (2017). Automated computer-based detection of encounter behaviours in groups of honeybees. Scientific Reports, 7(1), 1–9. 10.1038/s41598-017-17863-4.

10. Bozek, K., Hebert, L., Portugal, Y., Mikheyev, A. S., & Stephens, G. J. (2021). Markerless tracking of an entire honey bee colony. Nature Communications, 12(1), 1733. 10.1038/s41467-021-21769-1.

11. Cappeliez, P., & Moore, E. (1988). Effects of lithium on latent inhibition in the rat. Progress in Neuropychopharmacology & Biological Psychiatry, 12(4), 431–443. 10.1016/0278-5846(88)90103-0.

12. Castillo-Quan, J. I., Li, L., Kinghorn, K. J., Ivanov, D. K., Tain, L. S., Slack, C., Kerr, F., Nespital, T., Thornton, J., Hardy, J., Bjedov, I., & Partridge, L. (2016). Lithium Promotes Longevity through GSK3/NRF2-Dependent Hormesis. Cell Reports, 15(3), 638–650. 10.1016/j.celrep.2016.03.041.

13. Claudio, E. P., Rodriguez-Cruz, Y., Arslan, O. C., Giray, T., Agosto Rivera, J. L., Kence, M., Wells, H., & Abramson, C. I. (2018). Appetitive reversal learning differences of two honey bee subspecies with different foraging behaviors. PeerJ, 6, e5918. 10.7717/peerj.5918.

14. Core, R. T. (2020). Team. R: a language and environment for statistical computing. https://www.R-project.org/.

15. Costa, V. D., Tran, V. L., Turchi, J., & Averbeck, B. B. (2015). Reversal learning and dopamine: A Bayesian perspective. Journal of Neuroscience, 35(6), 2407–2416. 10.1523/JNEUROSCI.1989-14.2015

16. Davoudian, P. A., & Nitabach, M. N. (2021). Dopaminergic mechanism underlying reward encoding of punishment omission during reversal learning in Drosophila. Nature Communications. 10.1038/s41467-021-21388-w.

17. Dinges, C., Avalos, A., Abramson, C., Craig, D., Austin, Z., Varnon, C. A., … & Wells, H. (2013). Aversive conditioning in honey bees (Apis mellifera anatoliaca): a comparison of drones and workers. Journal of Experimental Biology, 216(21), 4124–4134. 10.1242/jeb.090100.

18. Erdem, B., Arslan, O. C., Sevin, S., Gozen, A. G., Agosto, J. L., Giray, T., & Alemdar, H. (2023). Effects of lithium on locomotor activity and circadian rhythm of honey bees. Scientific Reports, 13, 19861. 10.1038/s41598-023-46777-7.

19. Ferguson, H. J., Cobey, S., & Smith, B. H. (2001). Sensitivity to a change in reward is heritable in the honeybee, Apis mellifera. Animal Behaviour, 61(3), 527–534. 10.1006/anbe.2000.1635.

20. Franciscovich, A. L., Vrailas Mortimer, A. D., Freeman, A. A., Gu, J., & Sanyal, S. (2008). Overexpression screen in drosophila identifies neuronal roles of GSK-3β/shaggy as a regulator of AP-1-dependent developmental plasticity. Genetics, 180(4), 2057–2071. 10.1534/genetics.107.085555.

21. Geffre, A. C., Gernat, T., Harwood, G. P., Jones, B. M., Morselli Gysi, D., Hamilton, A. R., … & Dolezal, A. G. (2020). Honey bee virus causes context-dependent changes in host social behavior. PNAS, 117(19), 10406–10413. 10.1073/pnas.2002268117.

22. Giurfa, M. (2007). Behavioral and neural analysis of associative learning in the honeybee: A taste from the magic well. Journal of Comparative Physiology A: Neuroethology, Sensory, Neural, and Behavioral Physiology, 193(8), 801–824. 10.1007/s00359-007-0235-9.

23. Grigor’yan, G. A. (2014). The Role of Glycogen Synthase Kinase 3 in the Mechanisms of Learning and Memory. Neuroscience and Behavioral Physiology, 44(9), 1051–1058. 10.1007/s11055-014-0023-2.

24. Hammer, M., & Menzel, R. (1995). Learning and memory in the honeybee. The Journal of Neuroscience, 15(3), 1617–1630. 10.1523/JNEUROSCI.15-03-01617.1995.

25. Hedgepeth, C. M., Conrad, L. J., Zhang, J., Huang, H. C., Lee, V. M. Y., & Klein, P. S. (1997). Activation of the Wnt signaling pathway: A molecular mechanism for lithium action. Developmental Biology, 185(1), 82–91. 10.1006/dbio.1997.8552.

26. Hines, G., & Poling, T.H. (1984). Lithium effects on active and passive avoidance behavior in the rat. Psychopharmacology (Berl), 82, 78–82. 10.1007/BF00426385.

27. Hong, M., Chen, D. C. R., Klein, P. S., & Lee, V. M. Y. (1997). Lithium reduces tau phosphorylation by inhibition of glycogen synthase kinase-3. Journal of Biological Chemistry, 272(40), 25326–25332. 10.1074/jbc.272.40.25326.

28. Hooper, C., Markevich, V., Plattner, F., Killick, R., Schofield, E., Engel, T., Hernandez, F., Anderton, B., Rosenblum, K., Bliss, T., Cooke, S. F., Avila, J., Lucas, J. J., Giese, K. P., Stephenson, J., & Lovestone, S. (2007). Glycogen synthase kinase-3 inhibition is integral to long-term potentiation. European Journal of Neuroscience, 25(1), 81–86. 10.1111/j.1460-9568.2006.05245.x.

29. Hurst, V., Stevenson, P. C., & Wright, G. A. (2014). Toxins induce ’malaise’ behavior in the honeybee (Apis mellifera). Journal of Comparative Physiology A: Neuroethology, Sensory, Neural, and Behavioral Physiology, 200(10), 881–890. 10.1007/s00359-014-0932-0.

30. Ings, T. C., & Chittka, L. (2008). Speed-accuracy tradeoffs and false alarms in bee responses to cryptic predators. Current Biology, 18(19), 1520–1524. 10.1016/j.cub.2008.07.074.

31. Izquierdo, A., Brigman, J. L., Radke, A. K., Rudebeck, P. H., & Holmes, A. (2017). The neural basis of reversal learning: an updated perspective. Neuroscience, 345, 12–26. 10.1016/j.neuroscience.2016.03.021.

32. Jarriault, D., Fuller, J., Hyland, B. I., & Mercer, A. R. (2018). Dopamine release in mushroom bodies of the honey bee (Apis mellifera L.) in response to aversive stimulation. Scientific Reports, 8(1), 1–12. 10.1038/s41598-018-34460-1.

33. Jope, R. S. (2003). Lithium and GSK-3: One inhibitor, two inhibitory actions, multiple outcomes. Trends in Pharmacological Sciences, 24(9), 441–443. 10.1016/S0165-6147(03)00206-2.

34. Jope, R. S., Cheng, Y., Lowell, J. A., Worthen, R. J., Sitbon, Y. H., & Beurel, E. (2017). Stressed and Inflamed, Can GSK3 Be Blamed? Trends in Biochemical Sciences, 42(3), 180–192. 10.1016/j.tibs.2016.10.009.

35. Jovanovic, N. M., Glavinic, U., Ristanic, M., Vejnovic, B., Stevanovic, J., Cosic, M., & Stanimirovic, Z. (2022). Contact varroacidal efficacy of lithium citrate and its influence on viral loads, immune parameters and oxidative stress of honey bees in a field experiment. Frontiers in Physiology, 13, 1000944. 10.3389/fphys.2022.1000944.

36. Judd, L. L., Hubbard, B., Janowsky, D. S., Huey, L. Y., & Takahashi, K. I. (1977). The effect of lithium carbonate on the cognitive functions of normal subjects. Archives of General. Psychiatry, 34(3), 355–357. doi:10.1001/archpsyc.1977.01770150113013.

37. Kabra, M., Robie, A. A., Rivera-Alba, M., Branson, S., & Branson, K. (2013). JAABA: interactive machine learning for automatic annotation of animal behavior. Nature Methods, 10(1), 64–67. 10.1038/nmeth.2281.

38. Klein, P. S., & Melton, D. A. (1996). A molecular mechanism for the effect of lithium on development. Proceedings of the National Academy of Sciences of the United States of America, 93(16), 8455–8459. 10.1073/pnas.93.16.8455.

39. Kolics, É., Specziár, A., Taller, J., Mátyás, K. K. & Kolics, B. (2021a). Lithium chloride outperformed oxalic acid sublimation in a preliminary experiment for varroa mite control in pre wintering honey bee colonies. Acta Veterinaria Hungarica, 68(4), 370–373. 10.1556/004.2020.00060.

40. Kolics, É., Sajtos, Z., Mátyás, K., Szepesi, K., Solti, I., Németh, G., Taller, J., Baranyai, E., Specziár, A., & Kolics, B (2021b). Changes in lithium levels in bees and their products following anti-varroa treatment. Insects, 12(7), 579. 10.3390/insects12070579.

41. Kropf, D., & Müller-Oerlinghausen, B. (1979). Changes in learning, memory, and mood during lithium treatment. Acta Psychiatrica Scandinavica, 59(1), 97–124. 10.1111/j.1600-0447.1979.tb06951.x.

42. Menzel, R., & Müller, U. (1996). Learning and memory in honeybees: From behavior to neural substrates. Annual Review of Neuroscience, 19, 379–404. 10.1146/annurev.ne.19.030196.002115

43. Ngo, T. N.,Wu, K. C., Yang, E. C., Lin, T. T. (2019). A real-time imaging system for multiple honey bee tracking and activity monitoring. Computers and Electronics in Agriculture, 163, 104841. 10.1016/j.compag.2019.05.050.

44. Ngo, T. N., Rustia, D. J. A., Yang, E. C., Lin, T. T. (2021). Automated monitoring and analyses of honey bee pollen foraging behavior using a deep learning-based imaging system. Computers and Electronics in Agriculture, 187, 106239. 10.1016/j.compag.2021.106239.

45. Rein, C., Makosch, M., Renz, J. & Rosenkranz, P. (2022). Lithium chloride leads to concentration dependent brood damages in honey bee hives (Apis mellifera) during control of the mite Varroa destructor. Apidologie, 53, 38. 10.1007/s13592-022-00949-y.

46. Richter-Levin, G., Markram, H. & Segal, M. (1992). Spontaneous recovery of deficits in spatial memory and cholinergic potentiation of NMDA in CA1 neurons during chronic lithium treatment. Hippocampus, 2(3), 279–286. 10.1002/hipo.450020307.

47. Seeley, T. D. (1994). Honey bee foragers as sensory units of their colonies. Behavioral Ecology and Sociobiology, 34(1), 51–62. 10.1007/s002650050018.

48. Sevin, S., Bommuraj, V., Chen, Y., Afik, O., Zarchin, S., Barel, S., Arslan, O. C., Erdem, B., Tutun, H., & Shimshoni, J. A. (2022). Lithium salts: assessment of their chronic and acute toxicities to honey bees and their anti-Varroa field efficacy. Pest Management Science. 10.1002/ps.7071.

49. Shaldubina, A., Agam, G., & Belmaker, R. H. (2001). The mechanism of lithium action: State of the art, ten years later. Progress in Neuro-Psychopharmacology and Biological Psychiatry, 25(4), 855–866. 10.1016/S0278-5846(01)00154-3.

50. Siefert, P., Hota, R., Ramesh, V., & Grünewald, B. (2020). Chronic within-hive video recordings detect altered nursing behaviour and retarded larval development of neonicotinoid treated honey bees. Scientific Reports, 10(1), 1–15. 10.1038/s41598-020-65425-y.

51. Stambolic, V., Ruel, L., & Woodgett, J. R. (1996). Lithium inhibits glycogen synthase kinase-3 activity and mimics wingless signalling in intact cells. Current Biology, 6(12), 1664–1669. 10.1016/s0960-9822(02)70790-2.

52. Storozheva, Z. I., Gruden, M. A., Proshin, A. T., & Sewell, R. D. E. (2015). Learning ability is a key outcome determinant of GSK-3 inhibition on visuospatial memory in rats. Journal of Psychopharmacology, 29(7), 822–835. 10.1177/0269881115573805.

53. Tsaltas, E., Kontis, D., Boulougouris, V., Papakosta, V. M., Giannou, H., Poulopoulou, C., & Soldatos, C. (2007). Enhancing effects of chronic lithium on memory in the rat. Behavioural Brain Research, 177, 51–60. 10.1016/j.bbr.2006.11.003.

54. Vasconcellos, A.P. S., Tabajara, A.S., Ferrari, C., Rocha, E., & Dalmaz, C. (2003). Effect of chronic stress on spatial memory in rats is attenuated by lithium treatment. Physiology & Behavior. 79, 143–149. 10.1016/S0031-9384(03)00113-6.

55. Vergoz, V., Roussel, E., Sandoz, J. C., & Giurfa, M. (2007). Aversive learning in honeybees revealed by the olfactory conditioning of the sting extension reflex. PLoS ONE, 2(3). 10.1371/journal.pone.0000288.

56. Williams, R. S. B., & Harwood, A. J. (2000). Lithium therapy and signal transduction. Trends in Pharmacological Sciences, 21(2), 61–64. 10.1016/S0165-6147(99)01428-5.

57. Wolf, F. W., Eddison, M., Lee, S., Cho, W., & Heberlein, U. (2007). GSK-3/Shaggy regulates olfactory habituation in Drosophila. Proceedings of the National Academy of Sciences of the United States of America, 104(11), 4653–4657. 10.1073/pnas.0700493104.

58. Wu, C. L., Huang, L. T., Liou, C. W., Wang, T. J., Tung, Y. R., Hsu, H. Y., & Lai, M. C. (2001). Lithium-pilocarpine-induced status epilepticus in immature rats result in long-term deficits in spatial learning and hippocampal cell loss. Neuroscience Letters, 312(2), 113–117. 10.1016/S0304-3940(01)02202-9.

59. Xia, S., Liu, L., Feng, C., & Guo, A. (1997). Drug disruption of short-term memory in Drosophila melanogaster. Pharmacology Biochemistry and Behavior, 58(3), 727–735. 10.1016/S0091-3057(97)00045-2.

60. Zhan, F., Phiel, C. J., Spece, L., Gurvich, N., & Klein, P. S. (2003). Inhibitory phosphorylation of glycogen synthase kinase-3 (GSK-3) in response to lithium: Evidence for autoregulation of GSK-3. Journal of Biological Chemistry, 278(35), 33067–33077. 10.1074/jbc.M212635200.

61. Zhou, M., Chen, N., Tian, J., Zeng, J., Zhang, Y., Zhang, X., Guo, J., Sun, J., Li, Y., Guo, A., & Li, Y. (2019). Suppression of GABAergic neurons through D2-like receptor secures efficient conditioning in Drosophila aversive olfactory learning. Proceedings of the National Academy of Sciences of the United States of America, 116(11), 5118–5125. 10.1073/pnas.1812342116.

62. Ziegelmann, B., Abele, E., Hannus, S., Beitzinger, M., Berg, S., & Rosenkranz, P. (2018). Lithium chloride effectively kills the honey bee parasite Varroa destructor by a systemic mode of action. Scientific Reports, 8(1), 1–9; 10.1038/s41598-017-19137-5.

