## Supplementary Material for "Api-TRACE: A System for Honey Bee Tracking in a Constrained Environment to Study Bee Learning Process and the Effect of Lithium on Learning"

**Supplement Tables**

Table S1. Descriptive statistics for ESA assay.

|  | **Dose** | **n** | **mean** | **sd** | **se** | **med** | **quant25** | **quant75** | **IQR** | **min** | **max** |
| --- | --- | --- | --- | --- | --- | --- | --- | --- | --- | --- | --- |
| **Acquisition Phase** | Control | 27 | 138.799 | 45.807 | 8.816 | 140.700 | 115.333 | 156.350 | 41.017 | 34.967 | 289.000 |
|  | 5 mM | 33 | 131.839 | 38.417 | 6.687 | 138.567 | 114.833 | 156.367 | 41.533 | 20.467 | 192.700 |
|  | 25 mM | 31 | 143.492 | 19.993 | 3.591 | 147.200 | 141.767 | 150.650 | 8.883 | 55.400 | 180.133 |
|  | 125 mM | 28 | 141.633 | 34.656 | 6.549 | 148.950 | 129.550 | 160.567 | 31.017 | 52.200 | 211.900 |
| **Reversal Phase** | Control | 27 | 115.235 | 53.033 | 10.206 | 129.967 | 74.350 | 149.267 | 74.917 | 2.267 | 204.867 |
|  | 5 mM | 33 | 137.734 | 36.770 | 6.401 | 134.733 | 118.633 | 146.667 | 28.033 | 35.000 | 247.533 |
|  | 25 mM | 31 | 145.190 | 20.484 | 3.679 | 147.067 | 139.267 | 152.317 | 13.050 | 90.500 | 196.767 |
|  | 125 mM | 28 | 143.093 | 37.668 | 7.119 | 149.300 | 131.625 | 162.133 | 30.508 | 0.000 | 198.467 |

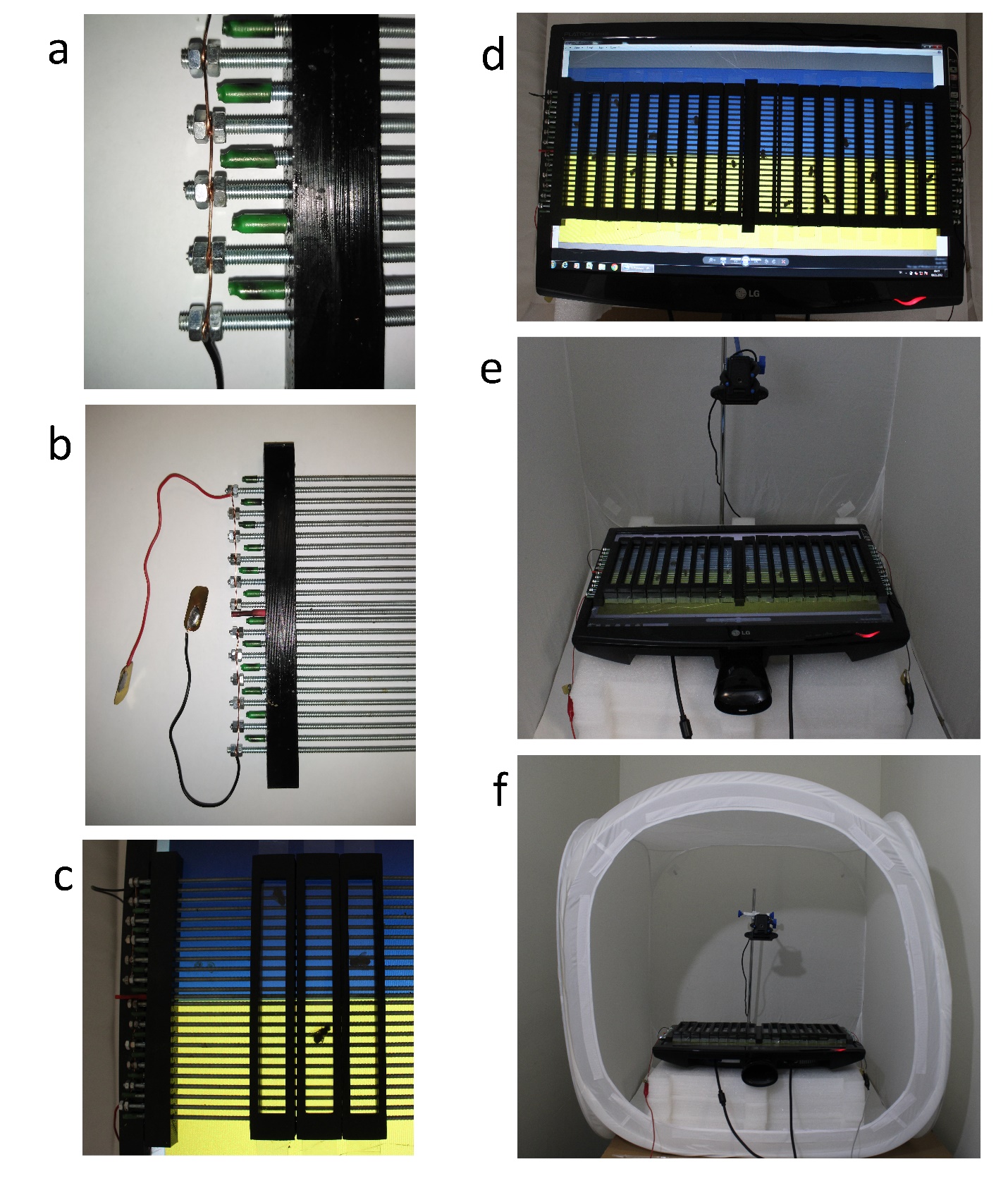

Figure S1. Experiment setup. Details of the electric grid (a, b) and experiment setup (c). The setup was positioned on the screen (d). All components were placed in a photo studio shooting tent (e). The experiment setup was fully prepared.
